# Sex-specific effects of voluntary wheel running on behavior and the gut microbiota-immune-brain axis in mice

**DOI:** 10.1101/2022.10.24.513258

**Authors:** Zoë AP Williams, Joanna Kasia Szyszkowicz, Natasha Osborne, Bshaier Allehyany, Christophe Nadon, Maryann Chinonye Udechukwu, Ana Santos, Marie-Claude Audet

**Author notes:** Correspondence to: Marie-Claude Audet, School of Nutrition Sciences, University of Ottawa, Ottawa, Ontario, K1H 8L1, Canada.

## Abstract

Physical exercise has been positioned as a promising strategy to prevent and/or alleviate anxiety and depression, but the mechanisms underlying its effects on mental health have yet to be entirely determined. Although the prevalence of depression and anxiety in women is about twice that of men, very few studies have examined whether physical exercise could affect mental health differently according to sex. This study examined, in mice, the sex-specific effects of voluntary exercise on body weight, depressive- and anxiety-like behaviors, as well as different markers along the gut microbiota-immune-brain axis. Male and female C57BL/6N mice had voluntary access to running wheels in their home-cages for 24 days or were left undisturbed in identical home-cages without running wheels. Behaviors were then examined in the open field, Splash, elevated plus maze, and tail suspension tests. Gene expression of pro-inflammatory cytokines, microglia activation-related genes, and tight junction proteins was determined in the jejunum and the hippocampus, while microbiota composition and predicted function were verified in cecum contents. Voluntary exercise limited weight gains, reduced anxiety-like behaviors, and altered grooming patterns in males exclusively. Although the exercise intervention resulted in changes to brain inflammatory activity and to cecal microbiota composition and inferred function in both sexes, reductions in the jejunal expression of pro-inflammatory markers were observed in females only. These findings support the view that voluntary exercise, even when performed during a short period, is beneficial for mental and intestinal health and that its sex-specific effects on behavior could be, at least in part, mediated by the gut microbiota-immune-brain axis.

## 1. Introduction

Depressive and anxiety disorders are leading causes of disability worldwide (Smith, 2014). Yet, available antidepressant and anxiolytic pharmacological treatments lead to complete remission in only about one third of individuals (Baldwin et al., 2011; Rush et al., 2006), highlighting the need to improve our understanding of alternative factors involved in the pathogenesis of these illnesses and to develop new targets for preventing and/or improving symptoms. Physical inactivity has been identified as one of the primary causes of chronic diseases (Małkiewicz et al., 2019) and associated with an increased risk and incidence of depression and anxiety (Gleeson et al., 2011; Pearce et al., 2022), suggesting that being physically inactive could be a predisposing factor to these disorders. In line with the possibility that exercise could limit anxiety and depressive symptoms, higher levels of physical activity have been associated with better mental health (Anderson & Shivakumar, 2013; De Moor et al., 2006; Ströhle, 2009). The antidepressant effects of exercise are also well-supported (Babyak et al., 2000; Blumenthal et al., 2007; Ignácio et al., 2019; Ross et al., 2022) and accumulating evidence points towards its anxiolytic properties (Asmundson et al., 2013; Wipfli et al., 2008).

The mechanisms involved in the antidepressant and anxiolytic effects of physical exercise have yet to be entirely determined. Emerging research suggests that the gut microbiota, the collection of microorganisms populating the gastrointestinal tract, may be part of a mechanistic system by which exercise improves mental health (Donoso et al., 2022; Gubert et al., 2020), potentially through its actions on the inflammatory signaling pathways between the gut environment and the brain (referred to as the gut microbiota-immune-brain axis). Individuals with depression and/or anxiety have a different gut microbiota (Chen et al., 2019; Kelly et al., 2016; Valles-Colomer et al., 2019; Zheng et al., 2016) and elevated levels of pro-inflammatory markers in the bloodstream and the brain compared to healthy individuals (Holmes et al., 2018; Hou et al., 2017; Osimo et al., 2020). A number of interventions targeting the gut microbiota have also been shown to reduce anxiety and depressive symptoms as well as circulating inflammatory factors in both healthy and clinical populations (Akkasheh et al., 2016; Hajifaraji et al., 2018; Messaoudi et al., 2011; Wallace & Milev, 2021). Supporting the possibility that physical activity could modulate the gut microbiota-immune-brain axis, exercise has been associated with changes in the gut microbiota (Campbell et al., 2016; O’Sullivan et al., 2015) and in intestinal permeability (Keirns et al., 2020; Luo et al., 2014), as well as in circulating levels of pro-inflammatory cytokines (Gonzalo-Encabo et al., 2021). In rodent models, exercise reduced basal levels of pro-inflammatory cytokines in intestinal lymphocytes (Hoffman-Goetz et al., 2009), limited stressor-induced microglial activation and cytokine elevations in the hippocampus (Xiao et al., 2021a), and prevented damages to the blood-brain barrier normally elicited by disease-based manipulations (Souza et al., 2017; Wolff et al., 2015).

Notably, although depression and anxiety are more prevalent in women (Baxter et al., 2013; Ferrari et al., 2013), most interventions aimed at reducing symptom severity and preventing and/or restoring biological impairments associated with these disorders have been developed in animal models using males and thus lack applicability to females (Rechlin et al., 2022). Among the studies that have looked at the psychological benefits of exercise, very few have considered sex as a biological variable in human populations, and most rodent studies only included males, thus adding to the underrepresentation of females in mental health prevention and treatment research. Notably, women and men with depression and anxiety differ in terms of gut microbiota and circulating inflammatory profiles, and in some cases these profiles are correlated with symptom severity in a sex-dependent manner (Audet, 2019). In rodent models of depression and/or anxiety, gut microbiota changes as well as circulating, intestinal, and brain inflammatory patterns, also differ between females and males (Bollinger et al., 2016; Wohleb et al., 2018), thus emphasizing the importance of examining these outcomes in both sexes.

To examine whether physical exercise limits the expression of behaviors suggestive of depression and anxiety in a sex-specific manner, and whether these effects are associated with changes to the gut microbiota-immune-brain axis, we provided home-cage running wheels to female and male mice for 24 days to give them voluntary access to exercise. We then assessed behaviors as well as inflammatory and tight junction factors in the jejunum, the primary site of nutrient and water absorption in the small intestine that harbors unique microbiota and immune systems (Bowcutt et al., 2014; Gu et al., 2013), and the hippocampus, a stress-sensitive brain region involved in depression (MacQueen & Frodl, 2011). Microbiota diversity, composition, and predicted metabolome were also determined from the contents of the cecum. With the knowledge that exercise influences the gut microbiota and has immunomodulatory properties, we hope that exploring the gut microbiota-immune-brain axis as a mechanism underlying the effects of physical activity on anxiety- and depressive-like behaviors will expand our understanding of exercise as a preventive and/or treatment approach for improving mental health. Furthermore, considering the role of sex as a biological variable in this regard will help optimize the efficacy of exercise interventions for mental health.

## 2. Materials and methods

### 2.1 Animals

Naïve female (*n*=20) and male (*n*=20) C57BL/6N mice (Charles River Laboratories, Montréal, QC), aged 9-10 weeks old, were singly housed in 39.6 × 21.5 × 17.2cm high-TEMP polysulfone cages (1285L Cage, Tecniplast) with a cardboard house, a cotton nestlet, and standard bedding. Mice had free access to food (Envigo 2014 Teklad global 14% protein rodent maintenance diet) and water and were maintained on a 12h light-dark cycle (lights on from 8:00h to 20:00h) with temperature and humidity kept between 21.0-23.0 °C and 30-50%, respectively. All experimental procedures were approved by the Animal Care Committee of the University of Ottawa (#3212), according to the guidelines of the Canadian Committee of Animal Care.

### 2.2 Summary of experimental procedures

Experimental procedures are summarized in Figure 1. After a few days of acclimation to their new environment, female and male mice were randomly assigned to an Exercise condition or a Control condition for 24 days (*n*/group = 10). Mice were weighed on Day 1 (Week 0) and then weekly on Day 8 (Week 1), Day 15 (Week 2), and Day 22 (Week 3). Behavioral testing occurred on Days 22 (open field test, Splash test) and 23 (elevated plus maze test, tail suspension test). All mice were euthanized on Day 24, approximately 16 hours after the last behavioral test, and brain (hippocampus) and gut (jejunum) sections as well as cecum contents were collected for the determination of gene expression of pro-inflammatory factors and tight junction proteins as well as of microbiota diversity, composition, and predicted metabolome. One female in the Exercise condition was removed from the study due to severe over-grooming behavior during the experiment.

**Figure 1:**
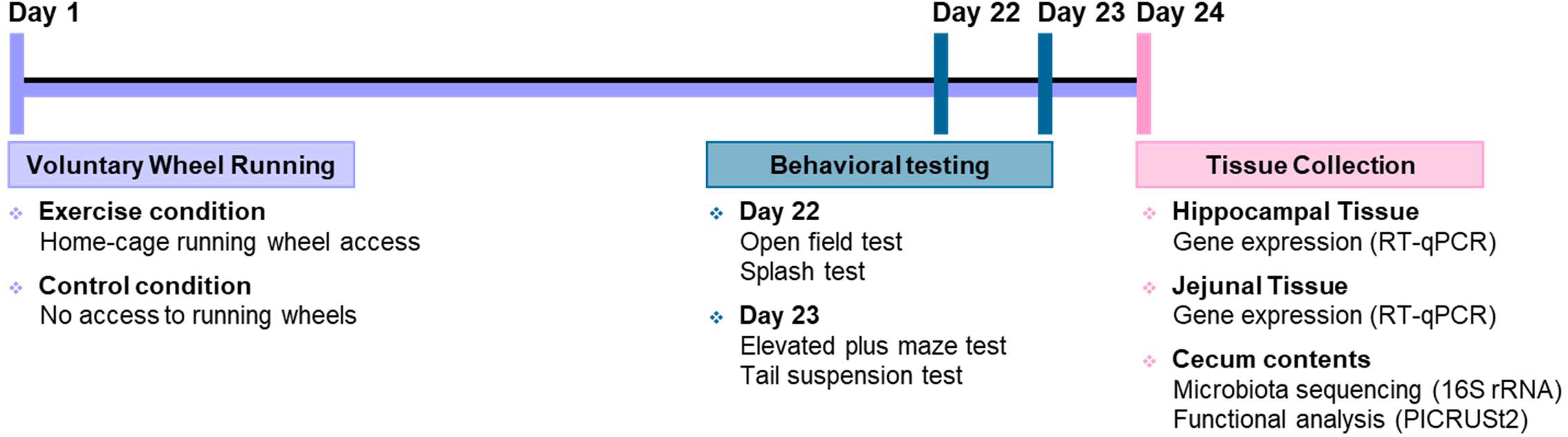
Summary of experimental procedures. Female (*n*=19) and male (*n*=20) C57BL/6N mice were given voluntary access to running wheels in their home-cages (Exercise condition) or were housed in identical cages without running wheels (Control condition) from Days 1 to 24. Body weights were measured on Day 1 (Week 0), before the introduction of running wheels into the home-cages or at corresponding times for mice in the Control condition, and then on Days 8 (Week1), 15 (Week 2), and 22 (Week 3). Behavior was assessed on Days 22 (open field and Splash tests) and 23 (elevated plus maze and tail suspension tests), and the hippocampus, the jejunum, and cecum contents were collected approximately 16 hours after on Day 24 for the determination of inflammatory and tight junction markers (RT-qPCR) and of microbiota diversity, composition, and inferred function (16S rRNA, PICRUST2).

### 2.3 Voluntary exercise procedures

Mice in the Exercise condition had voluntary access to home-cage running wheels from Days 1 to 24. Low-profile wireless running wheels (Med Associates Inc, ENV-047) were placed at one end of the home-cage, opposite to the food hopper, water bottle, and cardboard house. The numbers of wheel revolutions per minute were wirelessly transmitted from each running wheel to a hub connected to a laptop located in the housing room that recorded data using the Running Wheel Manager Data Acquisition Software (Med Associates Inc, SOF-736). The total distance traveled was calculated manually based on the total number of revolutions and the circumference traveled for each revolution, estimated from the external (15.5cm) and internal (9cm) diameter of the running wheels. Mice in the Control condition were housed in identical home-cages and housing conditions but without running wheels. On Days 22 and 23, during which behavioral testing occurred, running wheels remained in the home-cages but no data transmission and recording took place during daytime due to the intermittent presence of mice in the home-cages and the distance from the transmission hub. For this reason, the total running distance was determined from Days 1 to 22 only, specifically from the start of Day 1 light cycle to the end of Day 21 dark cycle.

### 2.4 Body weight

Mice were weighed before being placed into their respective experimental conditions on Week 0 (Day 1) and then once a week (Week 1 [Day 8], Week 2 [Day 15], and Week 3 [Day 22]). The percentage of weight change for each mouse from Week 0 was determined at the end of the exercise intervention as follows: [(Weight at a Week 3 - Weight at Week 0) / Weight at Week 0]*100.

### 2.5 Behavioral testing

#### 2.5.1 Open field test

The open field test was used to assess the innate fear of open and lit spaces, suggestive of anxiety-like behaviors, as previously described (Seibenhener & Wooten, 2015). Testing occurred on Day 22 under lighting standardized to approximately 300lx. The open field apparatus was composed of a 45 × 45 × 45cm arena made of opaque white acrylic plastic (Canus Plastics Incorporated). Mice were acclimatized to the testing room for 1 hour and then placed, individually, in the lower left corner of the open field and left to explore freely for 5 minutes. Between each trial, the apparatus was cleaned using a Quato 78 Plus solution. Movements were recorded using an overhead ceiling mounted camera and tracked with a video tracking software (Ethovision XT, version 11.5, Noldus). The time spent and the number of entries into regions defined as the center (15 × 15cm) and the four corners (each 10 × 10cm) of the field as well as the total distance traveled in the field were subsequently determined using Ethovision XT (version 11.5, Noldus).

#### 2.5.2 Splash test

The Splash test was used to assess motivational and self-care behaviors, suggestive of depressive-like behaviors (Isingrini et al., 2010). The test was administered on Day 22, around 2 hours following the open field test, under lighting standardized to approximately 530lx. Mice were acclimatized to the testing room 1 hour prior to the test. After gently spraying their lower back twice with water, mice were placed in an empty housing cage and grooming behavior was videotaped for 10 minutes. The total time spent grooming, mean length of grooming sessions, and total number of grooming sessions were subsequently scored using Observer XT (Noldus). Due to technical problems during the test, eight mice were excluded from the analyses and thus the sample size and degrees of freedom associated with Splash test parameters differ from the other behavioral outcomes.

#### 2.5.3 Elevated plus maze test

The elevated plus maze was used to assess the innate fear of open, lit, and elevated spaces, a feature of anxiety-like behaviors, as previously described (Lister, 1987; Walf & Frye, 2007). Testing occurred on Day 23 under lighting standardized to approximately 100lx. The apparatus, made of black acrylic plastic, was composed of one open arm (6-cm wide x 75-cm long) intersecting perpendicularly with one closed arm (6-cm wide x 75-cm long x 20-cm high) to form a cross shape. The entire structure was raised 74 cm above the floor. Mice were acclimatized to the testing room for 1 hour and then placed individually in the center area of the maze, facing the intersection of an open and a closed arm, and left to freely explore the apparatus for 5 minutes. Between each test trial, the apparatus was cleaned using Quato 78 Plus solution. Movements were tracked using an overhead ceiling mounted camera and a video tracking software (Ethovision XT, version 11.5, Noldus). Time spent in open and closed arms, number of entries into these arms, as well as the total distance traveled within the apparatus were subsequently calculated from the tracked movements using Ethovision XT (version 11.5, Noldus).

#### 2.5.4 Tail suspension test

The tail suspension test was used to assess passive coping behavior, a feature of depressive-like behaviors, as previously described (Cryan et al., 2005; Steru et al., 1985). The test occurred on Day 23, approximately 2 hours after the elevated plus maze test, under lighting standardized to around 100lx. The tail suspension apparatus was composed of a tail suspension interface cabinet (Med Associates Inc, DIG-735), a load cell amplifier hardware (Med Associates Inc, ENV-505TS), and an associated user interface software (Med Associates Inc, Tail Suspension SOF-821). Mice were acclimatized to the testing room 1 hour prior to the test, after which they were securely attached upside down to an aluminum bar by the tail with a small piece of surgical tape that prevented escape from the position. The aluminum bar was connected to a strain gauge and mice movements were recorded for 6 minutes. Time spent immobile was recorded by the strain gauge and the associated user interface software and used as the main measure of passive coping behavior. Between each test trial, the apparatus was cleaned using Quato 78 Plus solution. Two mice were excluded from the analysis due to technical problems during the test and thus the sample size and degrees of freedom associated with the time spent immobile differ from the other behavioral outcomes.

### 2.6 Euthanasia and tissue collection

On Day 24, approximately 16 hours after the last behavioral test, mice were euthanized by rapid decapitation. Whole brains were rapidly removed, placed on a stainless-steel brain matrix positioned on an ice plate (2.5 × 3.75 × 2.0cm; slots spaced approximately 500um apart), and sectioned coronally using razor blades guided by the matrix slots. The hippocampus was dissected from one of the brain sections based on the Franklin and Paxinos mouse atlas (Franklin & Paxinos, 2019). In parallel, the gastrointestinal tract was rapidly removed from the abdominal cavity and placed on a nuclease-free ice plate. From this, the jejunum was dissected after flushing its contents with a phosphate-buffered saline solution. As well, the contents from the cecum were collected. The hippocampus and jejunum sections, as well as contents collected from the cecum, were placed in nuclease-free cryotubes, flash frozen in liquid nitrogen, and stored at −80°C for the subsequent determination of pro-inflammatory and tight junction markers (hippocampus, jejunum) and microbiota parameters (cecum contents).

### 2.7 Reverse transcription-quantitative polymerase chain reaction (RT-qPCR)

Hippocampal and jejunal sections were homogenized using TRIzol and total RNA was extracted according to the manufacturer’s instructions (Invitrogen, Burlington, Canada). RNA concentration and purity were assessed using a NanoDrop™ One Spectrophotometer (ThermoFisher Scientific). Only RNA samples with 260/280 and 260/230 ratios between 1.80 and 2.20 were included and thus the sample size and degrees of freedom associated with hippocampal and jejunal gene expression differ from behavioral outcomes. Total RNA was then converted into cDNA via reverse transcription using iScript™ Reverse Transcription Supermix (Bio-Rad, Canada) and a T100 Thermal Cycler (Bio-Rad, Canada) and cDNA aliquots were analyzed in simultaneous real-time quantitative polymerase chain reactions using SsoAdvanced Universal SYBR Green Supermix (Bio-Rad, Canada) and a CFX96 Touch Real-Time PCR Detection System (Bio-Rad, Canada). Hippocampal and jejunal samples were analyzed in triplicates. Primers that amplified glyceraldehyde-3-phosphate dehydrogenase (GAPDH) and β-Actin (Actb) were used as reference genes and their geometric mean was used to normalize the expression of reference genes. Fold changes for the mRNA expression of genes of interest were calculated using the 2^−ΔΔCT^ method relative to the groups in the Control condition (females and males combined) (Livak & Schmittgen 2001, 2008). Primer sequences are listed in Supplementary Table 1.

### 2.8 Cecal microbiota analyses

#### 2.8.1 Genomic sequencing of the microbiota

DNA was extracted from cecum contents using a Stool Nucleic Acid Isolation Kit, according to manufacturer’s instructions (Norgen Biotek Corp, Thorold, Canada). Concentration and purity of DNA samples were assessed using a Qubit Fluorometer (ThermoFisher Scientific) with a few samples reassessed using Quant-iT™ PicoGreen (Invitrogen). The 16S ribosomal RNA (16S rRNA) gene amplicons were prepared according to Illumina 16S library preparation guidelines. Briefly, the V3 and V4 hypervariable regions of the 16S rRNA gene were amplified using the primers S-D-Bact-0341-b-S-17 (F: 5’ TCG TCG GCA GCG TCA GAT GTG TAT AAG AGA CAG CCT ACG GGN GGC WGC AG) and S-D-Bact-0785-a-A-21 (R: 5’ GTC TCG TGG GCT CGG AGA TGT GTA TAA GAG ACA GGA CTA CHV GGG TAT CTA ATC C) (Klindworth et al., 2013). The resulting amplicons were tagged with Illumina nucleotide sequencing adapters and dual-index barcodes for Illumina MiSeq compatibility and sample identification, respectively, after which the pooled library was sequenced on a MiSeq system using a 600-cycle MiSeq Reagent Kit v3 as per manufacturer’s instructions (Illumina, San Diego, CA, USA). Post-processing of the data resulting from the MiSeq sequencing was conducted using QIIME 2 (Bolyen et al., 2019), with paired-end reads passing a median quality score of Q ≥ 30 being kept and further denoised, filtered, and rarified using DADA2 (Callahan et al., 2016). Nearly all samples exceeded the retention threshold of 10,000 reads, except for one female from the Exercise condition with 8891 reads that was retained for completeness. The relative abundance of bacteria at various taxonomic levels was calculated for each sample following the alignment of reads to taxa using the Greengenes database (DeSantis et al., 2006). Preprocessed data was further analyzed using MicrobiomeAnalyst (Chong et al., 2020) to determine the Chao1 and Shannon alpha-diversity indices and the Bray-Curtis dissimilarity beta-diversity index. Betadiversity was calculated using Bray-Curtis distance and visualized using Principal Coordinate Analysis (PCoA). Permutational Multivariate Analysis of Variance (PERMANOVA) was used as the statistical method to confirm between-group differences.

#### 2.8.2 In silico prediction of microbiota metabolic capacity

The functional potential of the taxonomic changes was predicted using the Phylogenetic Investigation of Communities by Reconstruction of Unobserved States (PICRUSt2) Python package (Langille et al., 2013). In brief, the data generated from the 16S rRNA gene sequencing was used to infer the *in silico* bacterial metabolic capacity of each cecum sample. The abundance of Operational Taxonomic Units (OTUs) was processed using PICRUSt2 to predict the relative enrichment of the Kyoto Encyclopedia of Genes and Genomes (KEGG) orthologs, KEGG pathways, and their associated BRITE functional hierarchies (Kanehisa et al., 2012). Based on the *a priori* hypothesis that exercise would influence inflammatory-related pathways (Quiroga et al., 2020; Wang et al., 2020), KEGG pathways related to lipopolysaccharide biosynthesis, peptidoglycan biosynthesis, and NOD-like receptor signaling pathways were examined. Although this was not a primary aim of the study, because exercise is known to affect metabolism (Wang et al., 2020), as an exploratory measure we also conducted a set of analyses on KEGG pathways related to carbohydrate metabolism.

### 2.9 Statistical analyses

Data was analyzed using SPSS version 27 and graphed using GraphPad Prism version 9 (GraphPad Software Inc.). All data were first tested for normality (Shapiro-Wilk test) and homogeneity of variances (Levene test). Total distance traveled in running wheels, computed from the total number of revolutions, met normality and homogeneity variance assumptions and was analyzed using an unpaired t-test with Sex as the between-group factor (Females versus Males). Body weight over the course of the voluntary exercise procedure was analyzed using a 2 (Sex: Females versus Males) × 2 (Exercise: No exercise versus Voluntary wheel running) × 4 (Week: Weeks 0–3) mixed measures analyses of variance (ANOVA) with repeated measures on the within-group factor Week. Behavioral, hippocampal, and jejunal outcomes were compared using 2 (Sex: Females versus Males) x 2 (Exercise: No exercise versus Voluntary wheel running) between-group ANOVAs. Because of the multiple variables examined, microbiota-related data were compared using 2 (Sex) x 2 (Exercise) between-group multifactorial ANOVAs (MANOVAs). Follow-up comparisons of the main and simple effects of the ANOVAs and MANOVAs comprised *t* tests with a Bonferroni correction to maintain the family-wise error rate at 0.05. Relationships between the total distance run in the wheels (a measure of the total amount of exercise) and behavioral outcomes, hippocampal/jejunal inflammatory and tight junction marker fold changes, or cecal microbiota parameters changed by the exercise manipulation were determined using Pearson or Spearman correlation coefficients in each sex. Likewise, relationships between significant microbiota-related changes and behavioral outcomes were determined using Pearson or Spearman correlation coefficients in each sex. The alpha level was set to *p*<0.05 for all analyses, except for correlations where the alpha level was set at *p*<0.01 considering the higher number of variables considered.

## 3. Results

### 3.1 Comparable amount of voluntary exercise among females and males

We first determined whether females and males differed in the total amount of exercise performed on the running wheels. The total distance run in the wheels from Days 1 to 22, computed from the total number of revolutions completed during that period, was comparable among females and males, *t*_(17)_=1.207, *p*=.244 (Fig. 2).

**Figure 2:**
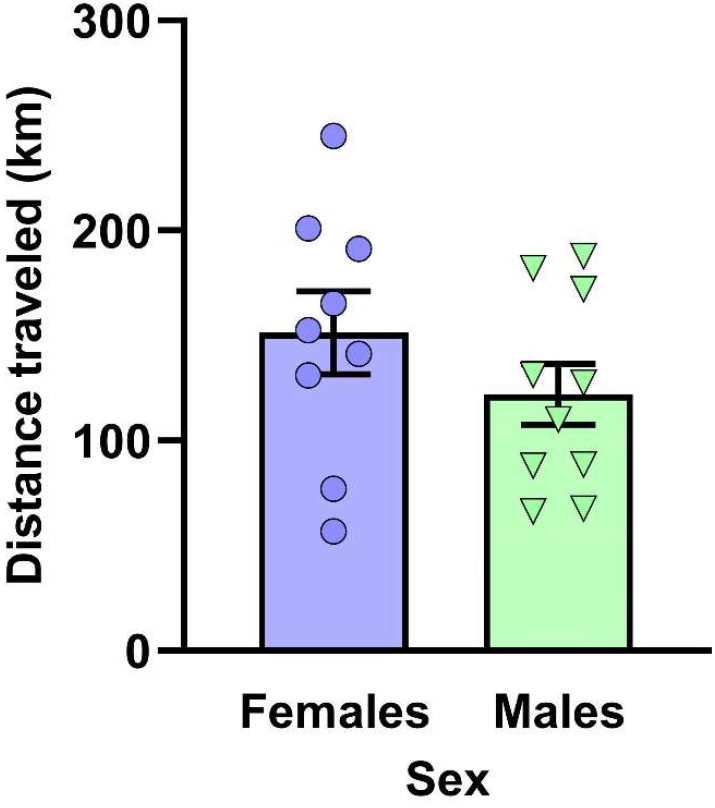
Female and male mice did the same amount of exercise during access to running wheels. The total distance run in the wheels (in km) from Days 1 to 22, computed form the total number of revolutions made during this period, was equivalent in female and male C57BL/6N mice with voluntary access to running wheels. Dots and triangles represent individual mice and bar plots and error bars represent group means ± S.E.M. (Females: *n*=9; males: *n*=10).

### 3.2 Voluntary access to exercise limited weight gains in males only

We next verified if having voluntary access to exercise influenced body weights and if this occurred in a sex-specific way. Body weights varied as a function of the interaction between Exercise, Sex, and Week, *F*_(3,105)_=5.145, *p*=.002, increasing over the weeks of the study, *F*_(3,105)_=42.812, *p*<.001, differently among males and females, *F*_(1,35)_=63.227, *p*<.001. As depicted in Figure 3A, follow-up comparisons of the simple effects comprising this interaction revealed that in males, body weights increased from Week 0 to Week 3 in those without access to running wheels (*p*<.001) but not in those that exercised (*p*=1.000). In females, body weights increased across the weeks of the study in both mice with and without exercise (*p*’s<.001). The percentages of weight change (weight lost or gained) from Week 0 to Week 3 also differed in mice that exercised, *F*_(1,35)_=7.523, *p*=.010, as well as in females versus males, *F*_(1,35)_=8.862, *p*=.005, and varied as a function of the Sex x Exercise interaction, *F*_(1,35)_=9.562, *p*=.004. Follow-up comparisons of the simple effects of this interaction confirmed that males that exercised gained less weight than males without exercise after Week 3 (*p*<.001), an effect that was not apparent in females (*p*=.809) (Fig. 3B).

**Figure 3:**
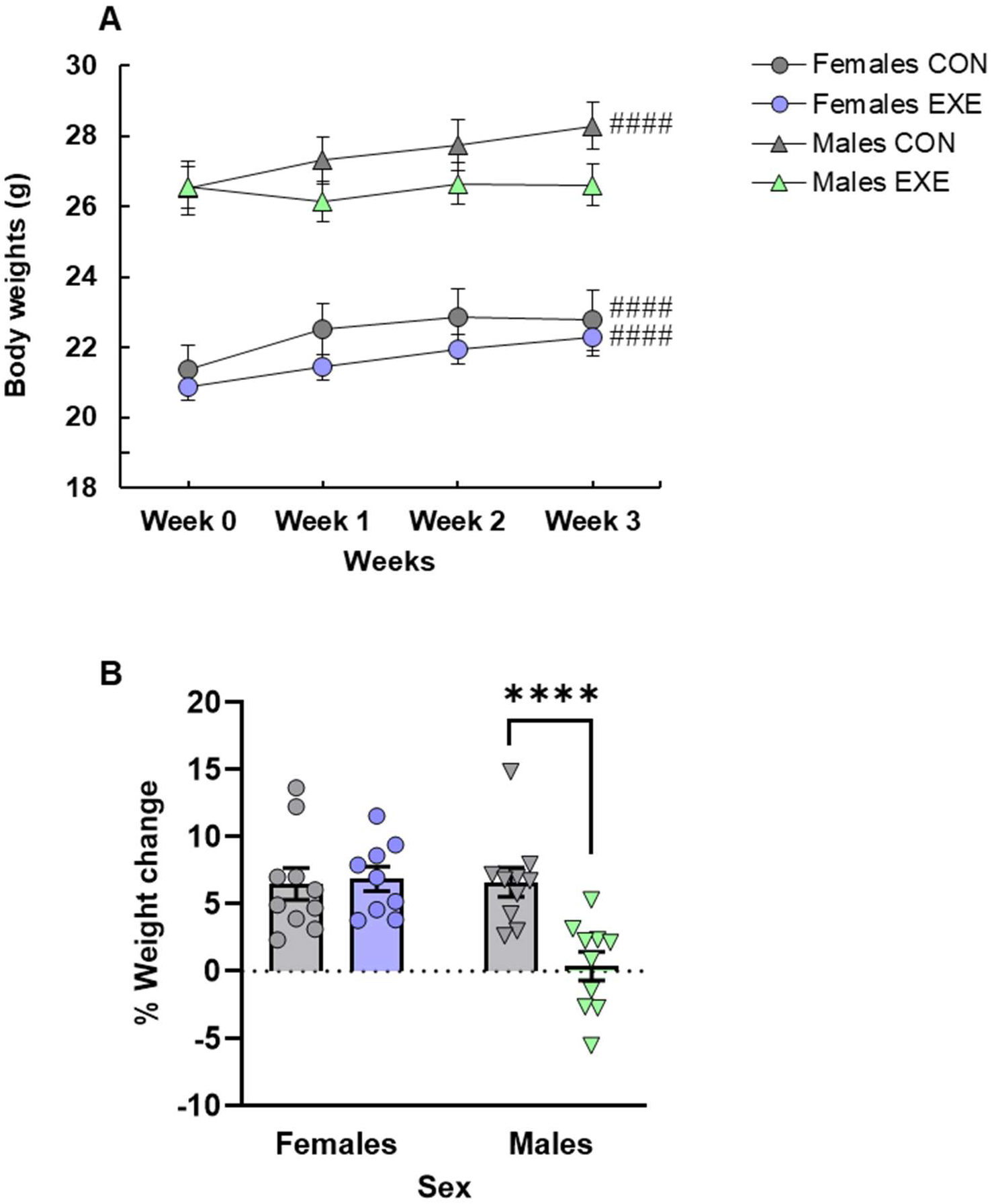
Access to running wheels prevented changes in body weight metrics in males only. (A) Actual weights from Week 0 (Day 1) to Week 3 (Day 22) increased in females with and without exercise as well as in males without exercise but remained unchanged in males with exercise. (B) Whereas females with exercise gained as much weight as their counterparts without exercise, males with exercise gained less weight than their counterparts without exercise. Dots and triangles represent individual mice and bar plots and error bars represent group means ± S.E.M. Females without exercise (Females CON: *n*=10); females with exercise (Females EXE: *n*=9); males without exercise (Males CON: *n*=10); males with exercise (Males EXE: *n*=10). ####*p*<.0001 relative to Week 0 and \*\*\*\**p*<.0001 relative to Males CON.

### 3.3 Behavior

#### 3.3.1 Open field test: voluntary access to exercise increased activity levels in both sexes

Although the standard indicators of fearful behaviors in the open field test remained unchanged by the different manipulations, mice that had access to exercise were more active in this environment, irrespective of sex (Supplementary Fig. 1). The time spent and the number of entries into the center of the field and the time spent in its corners were not affected by any of the manipulations (*p*’s>.05; Supplementary Fig. 1A-C). The number of entries into the corners, *F*_(1, 35)_=8.143, *p*=.007, and the distance traveled, *F*_(1,35)_=7.616, *p*=.009, however, were increased in exercised mice without being influenced by Sex or by the interaction between Sex and Exercise (*p*’s>.05; Supplementary Fig. 1D, E).

#### 3.3.2 Splash test: voluntary access to exercise changed grooming patterns in males only

As shown in Figure 4, male mice that exercised initiated more, although shorter, grooming sessions than males without exercise, while investing comparable time in grooming during the 10-minute test session, and these effects were not apparent in females. The total time engaged in grooming during the test was not affected by any of the manipulations (*p*’s>.05; Fig. 4A). In contrast, the number of grooming sessions, *F*_(1,27)_=6.671, *p*=.016, and the mean duration of these sessions, *F*_(1,27)_=6.254, *p*=.019, were affected by Exercise but not by Sex (*p*’s>.05). Although the interaction between Exercise and Sex for these factors did not reach significance (*p*’s>.05), based on the *a priori* predictions that behavioral outcomes in this test would be sexspecific, follow-up comparisons of the simple effects comprising this interaction were conducted. They confirmed that males that exercised engaged in grooming bouts more frequently (*p*=.010; Fig. 4B) and spent less time grooming during these bouts (*p*=.005; Fig. 4C) than males that did not exercise, but these effects were absent in females (*p*’s>.05).

**Figure 4:**
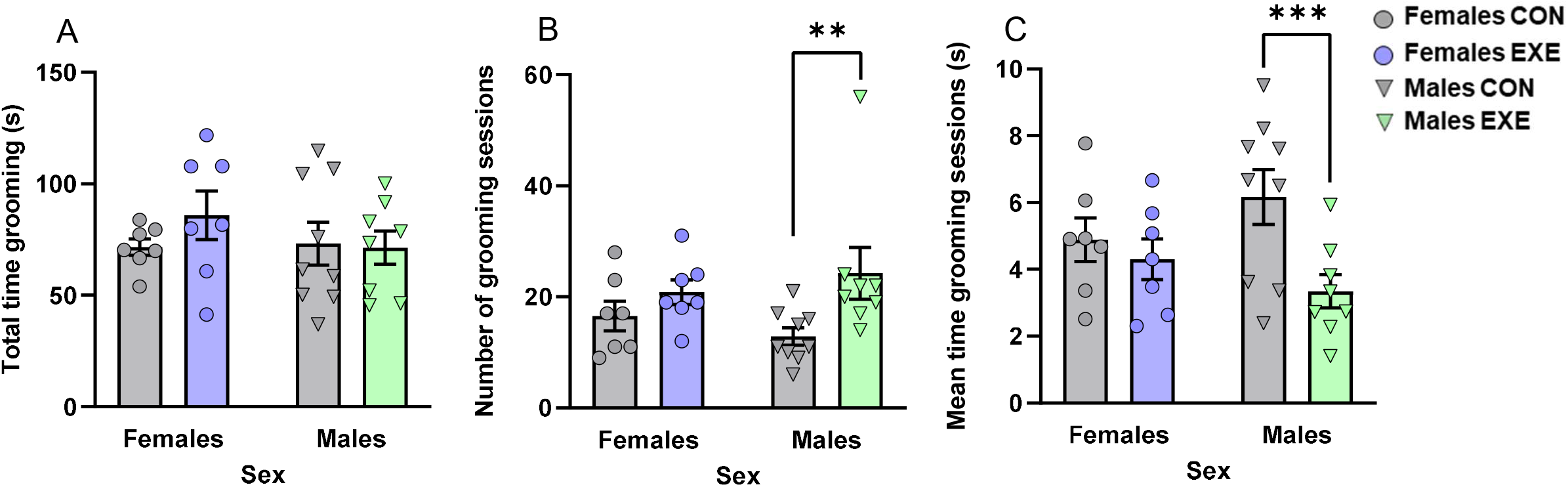
Access to running wheels changed grooming patterns in the Splash test in males only. (A) Total time spent grooming during the 10-min test session (in seconds [s]) was comparable among groups. (B) Males with exercise initiated more grooming sessions than those without exercise. (C) Males with exercise spent less time in average during these grooming sessions (in seconds [s]) than those without exercise. Dots and triangles represent individual mice and bar plots and error bars represent group means ± S.E.M. Females without exercise (Females CON: *n*=7); females with exercise (Females EXE: *n*=7); males without exercise (Males CON: *n*=9); males with exercise (Males EXE: *n*=8). \*\**p*<.01 and \*\*\**p*<.005 relative to Males CON.

#### 3.3.3 Elevated plus maze test: voluntary access to exercise reduced the fear of opened areas of the maze in males only

Voluntary access to exercise decreased the main indicators of fearful behaviors in the elevated plus maze in males but not in females (Fig. 5). The time in the open arms varied as a function of the Exercise x Sex interaction, *F*_(1,35)_=4.795, *p*=.035, and was influenced by both Exercise, *F*_(1,35)_=3.969, *p*=.054, and Sex, *F*_(1,35)_=6.479, *p*=.015. The time in the closed arms, although just shy of significance in terms of the Exercise x Sex interaction, *F*_(1,35)_=4.096, *p*=.051, was significantly affected by Sex, *F*_(1,35)_=10.537, *p*=.003, and moderately by Exercise, *F*_(1,35)_=3.411, *p*=.073. Follow-up comparisons of the simple effects comprising the Sex x Exercise interactions confirmed that male mice that exercised spent more time in the open arms (*p*=.005; Fig. 5A) and less time in the closed arms (*p*=.009; Fig. 5B) compared to males without exercise, an effect that was not apparent in females (*p*’s>.05). As well, both Exercise, *F*_(1,35)_=7.004, *p*=.012, and Sex, *F*_(1,35)_=4.326, *p*=.045, affected the number of entries into the open arms, but not entries into the closed arms (*p*’s>.05; Fig. 5D). Although the interaction between these two factors was not significant, *F*_(1,35)_=1.177, *p*=.285, based on the *a priori* predictions that behavioral outcomes in this test would be sex-dependent, follow-up comparisons of the simple effects comprising this interaction were conducted and revealed that males that exercised entered open arms more frequently than males without exercise (*p*=.011; Fig. 5C). Finally, the distance traveled in the maze was not affected by any of the manipulations (*p*’s>.05; Fig. 5E).

**Figure 5:**
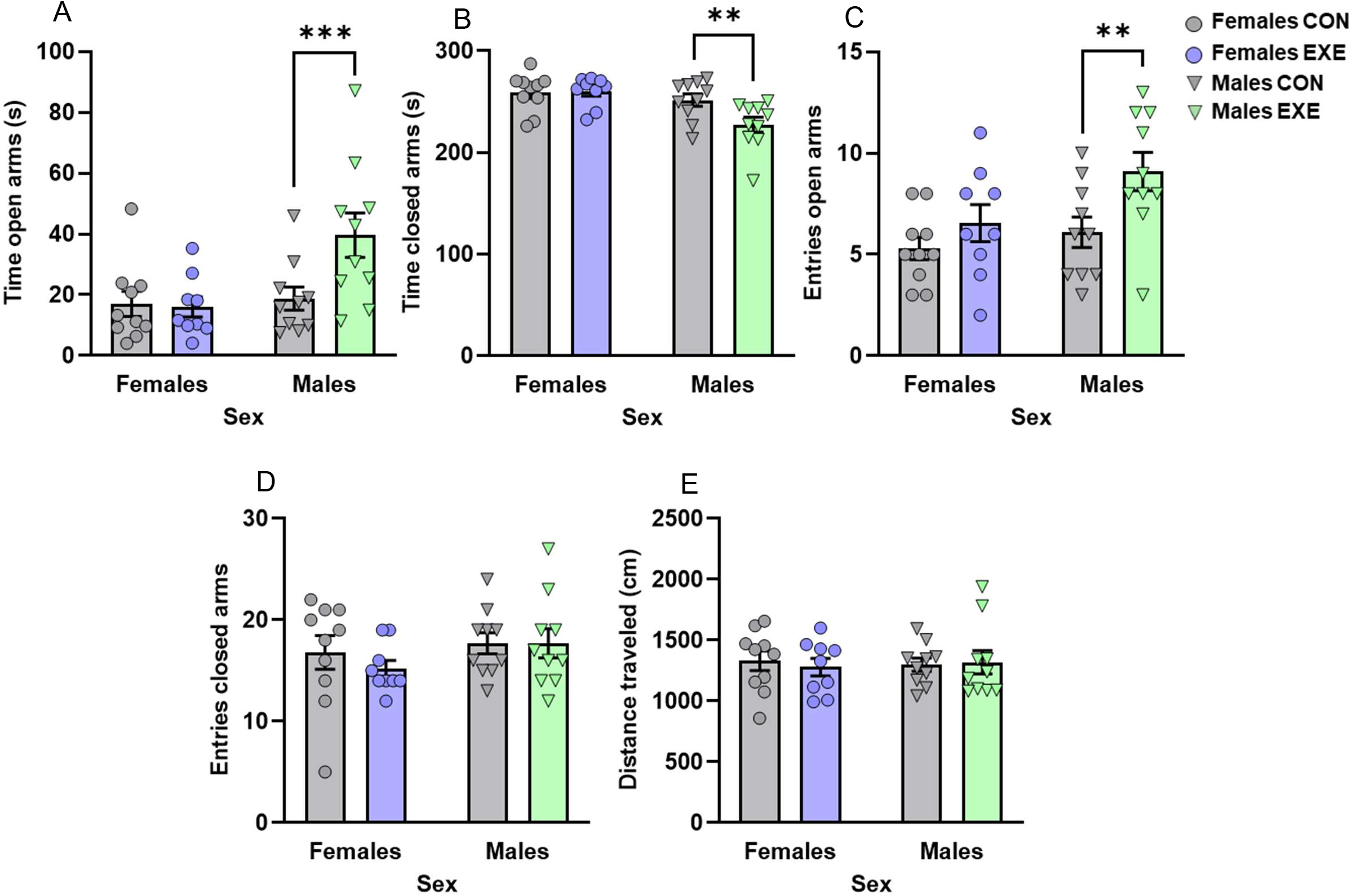
Access to running wheels reduced the fear of open arms in the elevated plus maze in males only. Males with exercise spent (A) more time in the open arms of the elevated plus maze (in seconds [s]) and (B) less time in the closed arms of the maze (in seconds [s]) than those without exercise. Males with exercise made (C) more entries into the open arms of the maze but (D) comparable entries into the closed arms than those without exercise. (E) The distance traveled in the apparatus (in cm) was comparable among groups. Dots and triangles represent individual mice and bar plots and error bars represent group means ± S.E.M. Females without exercise (Females CON: *n*=10); females with exercise (Females EXE: *n*=9); males without exercise (Males CON: *n*=10); males with exercise (Males EXE: *n*=10). \*\**p*<.01 and \*\*\**p*<.005 relative to Males CON.

#### 3.3.4 Tail suspension test: voluntary access to exercise had no effect on passive coping behavior

The time spent immobile in the tail suspension test was not affected by Sex, Exercise, or the interaction between these two factors (*p*’s>.05; Supplementary Fig. 1F).

### 3.4 Voluntary access to exercise downregulated CX3C chemokine receptor 1 (Cx3cr1) within the hippocampus in both sexes

Among all the genes targeted in the hippocampus, only Cx3cr1, the receptor for the chemokine fractalkine (Cx3cl1) expressed primarily by central nervous system resident microglia (Harrison et al., 1998), was changed by the manipulations, being lower in mice that exercised, *F*_(1,26)_=5.407, *p*=.028, irrespective of Sex and of its interaction with Exercise (*p*’s>.05) (Fig. 6A). Although the expression of brain-derived neurotrophic factor (BDNF) in the hippocampus appeared moderately increased in mice that exercised, the changes did not reach significance, *F*_(1,26)_=3.474, *p*=.074, and were not influenced by Sex and by its interaction with Exercise (*p*’s>.05; Fig. 6B). None of the pro-inflammatory cytokines or the tight junction proteins in the hippocampus was changed by Exercise or by Sex (*p*’s>.05; Supplementary Table 2).

**Figure 6:**
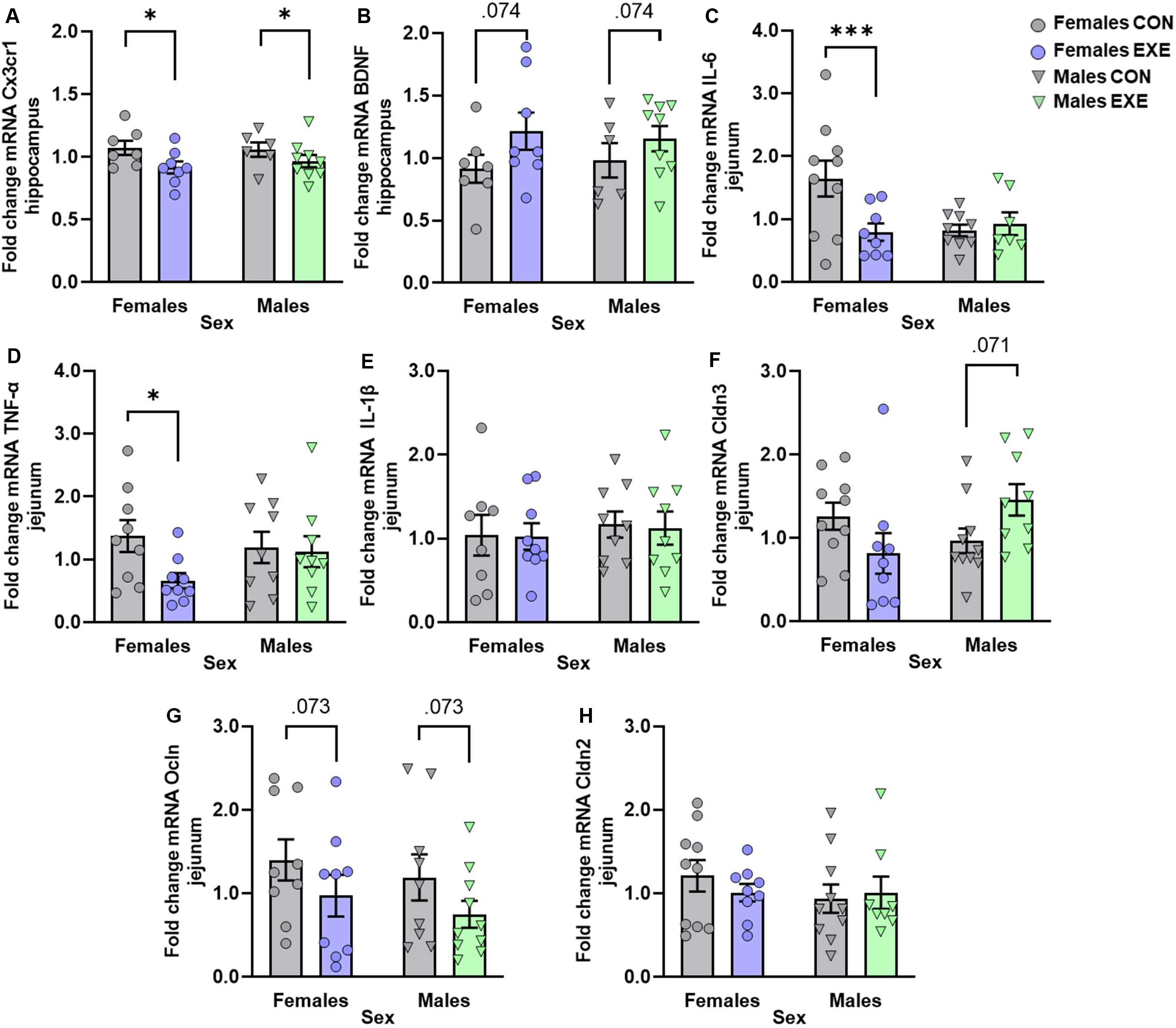
Gene expression of inflammatory and tight junction markers in the hippocampus and jejunum of female and male mice with or without access to running wheels. (A) Hippocampal expression of Cx3cr1 was reduced in both females and males with exercise whereas (B) the apparent hippocampal increases in BDNF expression in females and males with exercise did not reach significance. Females with exercise had lower jejunal expression of (C) IL-6 and of (D) TNF-α than their counterparts without exercise whereas (E) IL-1β was unchanged by any of the manipulations. The modest changes in (F) Cldn3 and (G) Ocln expression in the jejunum after exercise did not reach significance. (H) Jejunal Cldn2 expression was comparable among groups. Dots and triangles represent individual mice and bar plots and error bars represent group means ± S.E.M. Females without exercise (Females CON: *n*=7-10); females with exercise (Females EXE: *n*=8-9); males without exercise (Males CON: *n*=6-10); males with exercise (Males EXE: *n*=7-10). \**p*<.05 and \*\*\**p*<.005 relative to sex-matched CON.

### 3.5 Voluntary access to exercise downregulated jejunal pro-inflammatory cytokines in females only

Voluntary access to exercise reduced pro-inflammatory cytokine expression within the jejunum in females and had modest but non-significant effects on tight junction proteins across sexes (Fig. 6). Jejunal expression of interleukin (IL)-6 varied as a function of the Exercise x Sex interaction, *F*_(1,30)_=5.509, *p*=.026, with the apparent effects of Exercise, *F*_(1,30)_=3.328, *p*=.078, and Sex, *F*_(1,30)_=2.284, *p*=.099, being non-significant. Follow-up comparisons of the simple effects comprising this interaction confirmed that IL-6 was lower in females that exercised compared to females without exercise (*p*=.005; Fig. 6C). Although the main effects of Exercise and Sex and the interaction between the two factors did not reach significance for tumor necrosis factor (TNF)-α and IL-1β (*p*’s>.05), based on the *a priori* predictions that jejunal inflammatory status after exercise would be sex-specific, follow-up comparisons of the simple effects comprising this interaction were conducted and confirmed that exercise in females downregulated TNF-α (*p*=.032; Fig. 6D), but not IL-1β (*p*=.958; Fig. 6E). In relation to tight junction proteins, the expression of claudin-3 (Cldn3) varied as a function of the Exercise x Sex interaction, *F*_(1,34)_=6.315, *p*=.017, without being influenced by Exercise or Sex (*p*’s>.05). Follow-up comparisons of the simple effects comprising this interaction revealed that the apparent Cldn3 elevations in exercised males only approached significance (*p*=.071; Fig. 6F). Similarly, occludin (Ocln) reductions in exercised mice failed to reach significance, *F*_(1,33)_=3.438, *p*=.073 (Fig. 6G), irrespective of Sex and of its interaction with Exercise (*p*’s>.05). Finally, Cldn2 was not affected by any of the manipulations (*p*’s>.05; Fig. 6H).

### 3.6 Cecal microbiota

#### 3.6.1 Voluntary access to exercise decreased microbiota richness in males only

We first verified if voluntary access to exercise changed microbiota diversity metrics and if this occurred in a sex-dependent manner. The Chao1 index, reflective of bacterial species richness, varied as a function of the Exercise x Sex interaction, *F*_(1,35)_=4.759, *p*=. 036, without being directly affected by Exercise or Sex (*p*’s>.05). Follow-up comparisons of the simple effects comprising this interaction confirmed that Chao1 was lower in males with exercise compared to their counterparts without exercise (*p*=.016) and this was not seen in females (*p*=.573; Fig. 7A). In contrast, the Shannon index (Fig. 7B) and beta-diversity (Fig. 7C) were not affected by any of the manipulations (*p*’s>.05).

**Figure 7:**
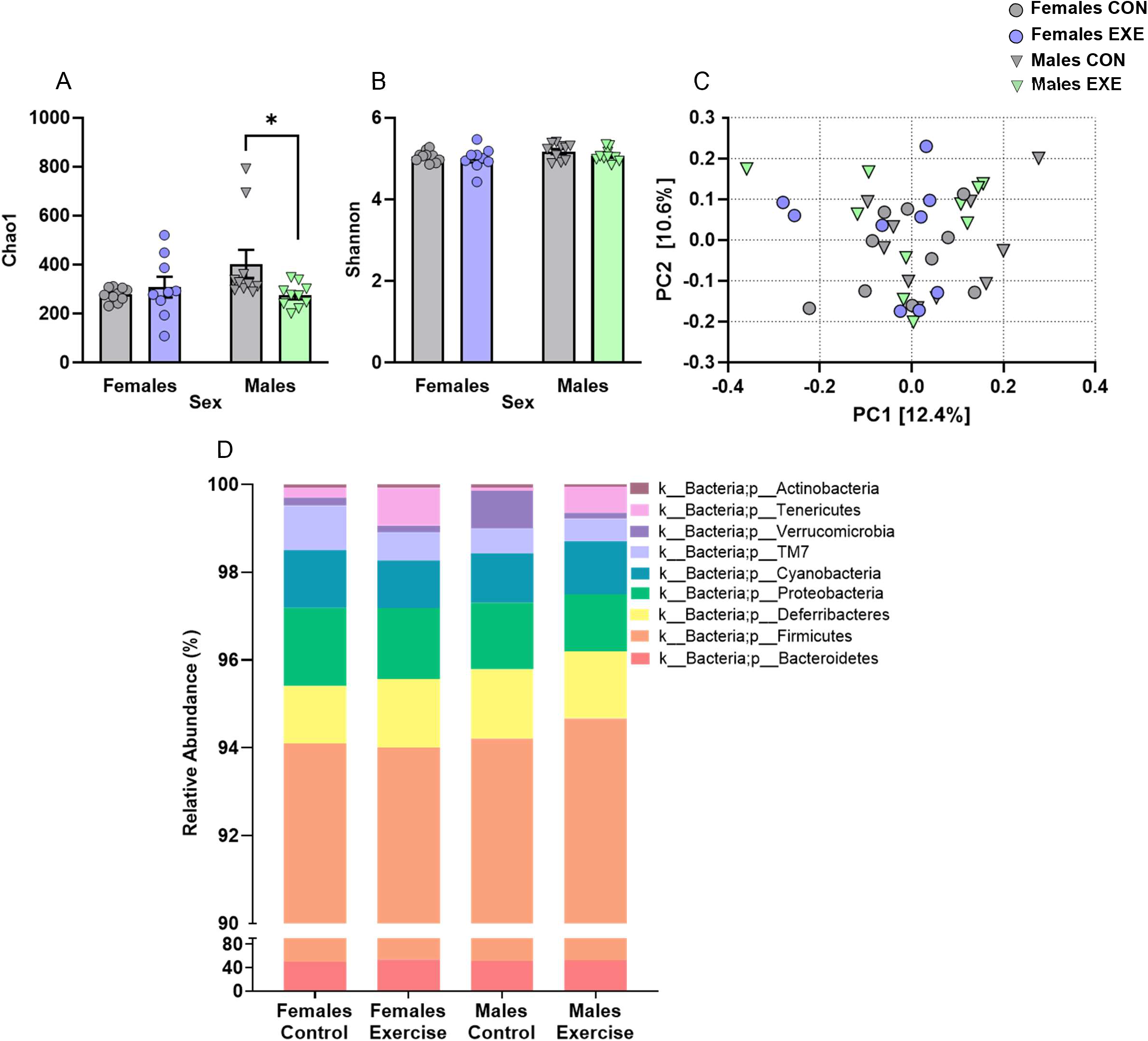
Alpha- and beta-diversity metrics and relative abundance of phyla in contents from the cecum of female and male mice with or without access to running wheels. (A) The Chao1 index of bacterial species richness was reduced in males with exercise compared to males without exercise whereas (B) the Shannon diversity index was comparable among groups. (C) Beta-diversity as illustrated by Bray-Curtis dissimilarity was comparable among all groups. (D) Relative abundance of phyla was unaltered by Exercise or Sex. Dots and triangles represent individual mice and bar plots and error bars represent group means ± S.E.M. Stacked bars represent the mean of the relative abundance of each phylum per group. Females without exercise (Females CON: *n*=10); females with exercise (Females EXE: *n*=9); males without exercise (Males CON: *n*=10); males with exercise (Males EXE: *n*=10). \**p*<.05 relative to Males CON.

#### 3.6.2 Voluntary access to exercise changed microbiota composition in cecum contents at the family and genus taxonomic levels in a sex-specific manner

Among the 9 phyla detected in the cecum contents of mice, none were affected by whether mice had voluntary access to exercise or by whether they were females or males (*p*’s>.05; Fig. 7D). Despite the absence of group differences at the phylum level, changes in the relative abundance of a number of families and genera specific to exercise and/or sex were observed within the phyla Firmicutes, Bacteroidetes, and Proteobacteria. Among the 67 families and 113 genera detected in cecum contents, 11 families and 25 genera were present in one sample only and thus were excluded from the subsequent analyses. In the taxa with higher abundance (>1%), only the *S24_7* family (Bacteroidetes phylum; accounting for 23.290% of the identified OTUs) differed among the experimental groups, being increased in mice that exercised, *F*_(1,35)_=4.517, *p*=.041, irrespective of Sex and its interaction with Exercise (*p*’s>.05; Fig. 8A). Similar to *S24_7*, the less abundant *Comamonadaceae* family (Proteobacteria phylum; 0.006% of the identified OTUs) was increased in exercised mice, *F*_(1,35)_=4.660, *p*=.038, irrespective of Sex and of its interaction with Exercise (*p*’s>.05; Fig. 8B). In contrast to changes specific to exercise, the *Lactonifactor* genus (Firmicutes phylum; 0.308% of the identified OTUs), *F*_(1,35)_=4.775, *p*=.036, the *Rhizobiaceae* family (Proteobacteria phylum, represented entirely by its genus *Rhizobium*; 0.042% of the identified OTUs), *F*_(1,35)_=4.583, *p*=.039, as well as the *Barnesiellaceae* family (Bacteroidetes phylum, represented entirely by its genus *Barnesiella*; 0.011% of the identified OTUs), *F*_(1,35)_=5.581, *p*=.024, differed between sexes only, with female mice having more *Lactonifactor* than males, but less *Rhizobium* and *Barnesiella* (Supplementary Fig. 2).

**Figure 8:**
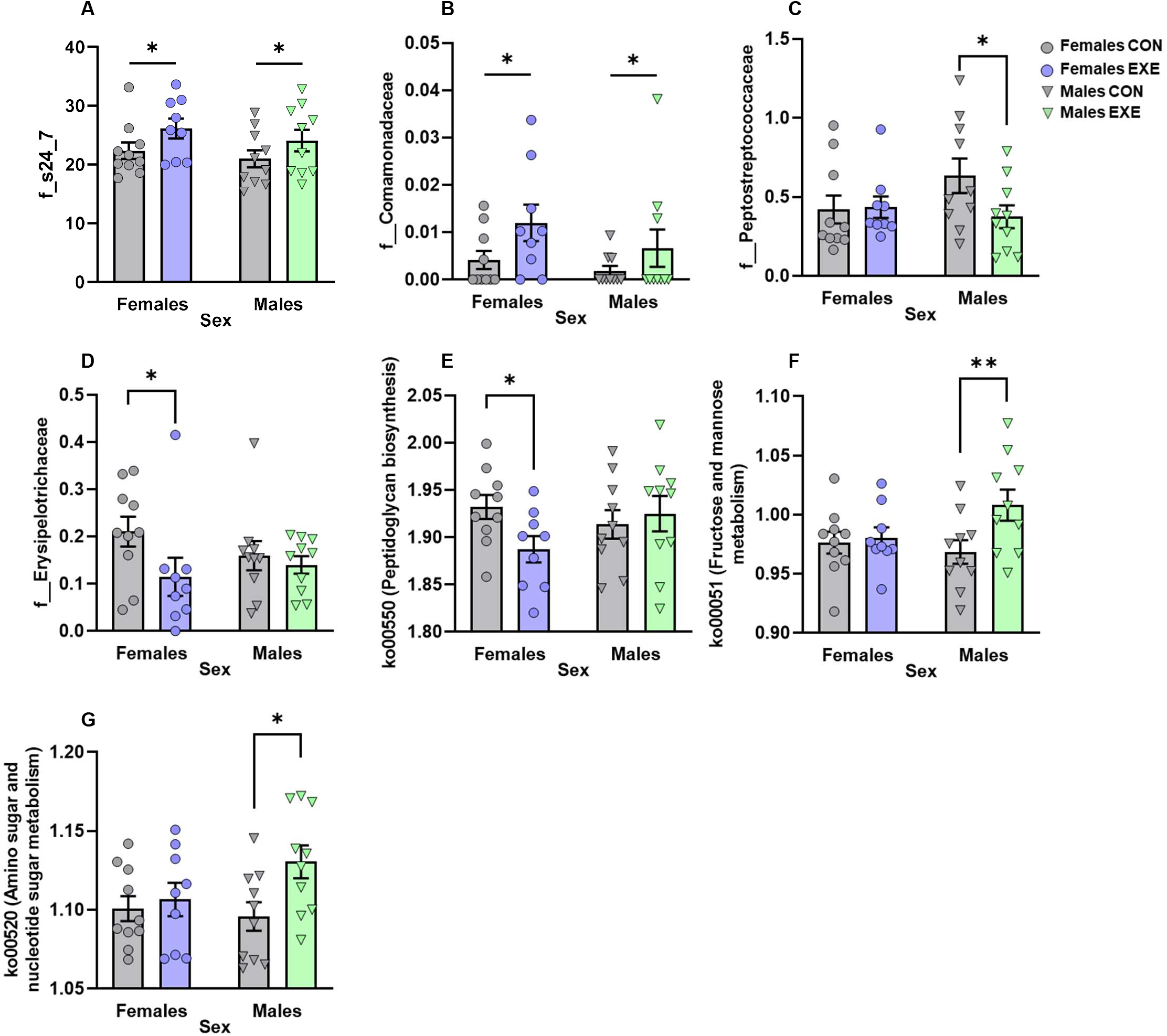
Relative abundance of families and genera in contents from the cecum of female and male mice with or without access to running wheels. Relative abundances of the *s24-7* (A) and *Comamonadaceae* (B) families were increased in both females and males with exercise. Access to exercise promoted sex-specific changes in the *Peptostreptococcaceae* and *Erysipelotrichaceae* families, with *Peptostreptococcaceae* being decreased in males (C) and *Erysipelotrichaceae* being decreased in females (D). Sex-dependent effects of exercise in KEGG pathways related to inflammation and carbohydrate metabolism, with the peptidoglycan biosynthesis pathways (ko00550) being decreased in females (E) and the fructose and mannose metabolism (ko00051; F) and amino sugar and nucleotide sugar metabolism (ko00520; G) being increased in males. Dots and triangles represent individual mice and bar plots and error bars represent group means ± S.E.M. Females without exercise (Females CON: *n*=10); females with exercise (Females EXE: *n*=9); males without exercise (Males CON: *n*=10); males with exercise (Males EXE: *n*=10). \**p*<.05 and \*\**p*<.01 relative to sex-matched CON.

A few taxonomic changes resulted from the exercise procedure in one sex only. Based on reports that *Peptostreptococcaceae* and *Erysipelotrichaceae* were modulated by exercise (Kaakoush, 2015; Li et al., 2021; Zhao et al., 2018; Zhu et al., 2020), the simple effects comprising the Sex x Exercise interaction for these two families were examined, despite no main effects or interaction being significant (*p*’s>.05). These analyses revealed that exercised males had less *Peptostreptococcaceae* (*p*=.041; Proteobacteria phylum, accounting for 0.467% of the identified OTUs; Fig. 8C) whereas exercised females had less *Erysipelotrichaceae* (*p*=.016; Firmicutes, accounting for 0.157% of the identified OTUs; Fig. 8D) compared to their sex-matched counterparts without access to exercise.

#### 3.6.3 Voluntary access to exercise changed the predicted metabolic capacity of cecal bacteria in a sex-dependent manner

Using PICRUSt2, based on abundance levels of OTUs obtained from the 16S rRNA gene sequencing data, we inferred the metabolic capacity of each cecal sample. Based on the *a priori* prediction that physical exercise would modulate inflammatory-related pathways (Quiroga et al., 2020; Wang et al., 2020), analyses were conducted on the KEGG pathways related to lipopolysaccharide biosynthesis, peptidoglycan biosynthesis, and NOD-like receptor signaling. From these, only peptidoglycan biosynthesis varied moderately as a function of the Exercise x Sex interaction *F*_(1,35)_=3.306, *p*=.078, without main effects of Sex or Exercise (*p*’s>.05). Followup comparisons of the simple effects comprising this interaction confirmed that exercise downregulated the peptidoglycan biosynthesis pathway in females (*p*=.051) but not in males (*p*=.600; Fig. 8E). As an exploratory measure, we also examined the effects of sex and exercise on KEGG pathways related to carbohydrate and lipid metabolism. From these, fructose and mannose metabolism (*F*_(1,35)_=4.366, *p*=.044) and amino sugar and nucleotide sugar metabolism (*F*_(1,35)_=4.515 *p*=.019) were influenced by Exercise. Although the interaction between Exercise and Sex was not significant (*p*’s>.05), based on the *a priori* prediction that physical exercise would modulate metabolism-related pathway in a sex-specific way, follow-up comparisons of the simple effects comprising this interaction were conducted. They confirmed that exercise upregulated the fructose and mannose metabolism and amino sugar and nucleotide sugar metabolism pathways in males (*p*’s=.010 and .013) but not in females (*p*’s>.05; Fig. 8F, G).

### 3.7 Amounts of exercise were correlated with microbiota changes in a sex-dependent manner

We verified whether behaviors and components of the gut microbiota-immune-brain axis that changed after voluntary running were related to how much mice exercised. None of the behaviors or of the jejunal and hippocampal targets that were significantly modified in mice with access to exercise were related to the total distance run in the wheels during this period (Supplementary Fig. 3). Although these relationships did not reach the alpha level of *p*<.01 set for significance, the relative abundance of *Peptostreptococcaceae* was positively correlated to the amounts of exercise in females, *r*_(9)_=0.683, *p*=.042, while that of the *Lactonifactor* genus was negatively correlated to amounts of exercise in males, *r*_(10)_=-0.660, *p*=.039 (Supplementary Fig. 3). None of the KEGG inflammatory- or metabolism-related pathways examined were linked to the amounts of exercise (*p*’s>.05).

### 3.8 Behavioral parameters changed by exercise were linked to specific taxa and KEGG pathways in a sex-dependent way

We last assessed whether behaviors suggestive of depressive- and anxiety-like phenotypes were related to cecal microbiota composition and predicted metabolic capacity in each sex separately (Fig. 9). Curiously, more significant relationships were detected in females than in males, despite the fact that behavioral outcomes in this sex had not been changed by exercise. The Chao1 index of bacterial species richness was positively correlated with the time spent in the open arms of the elevated plus maze, *r*_(19)_=0.660, *p*=.002, and with the number of entries in these arms, *r*_(19)_=0.602, *p*=.006, but negatively correlated with the time spent in the closed arms of the maze, *r*_(19)_=-0.637, *p*=.003. Similarly, *Barnesiella* abundance was positively correlated with the time spent in the open arms of the elevated plus maze, *r*_(19)_=0.593, *p*=.007. Finally, enrichment of the peptidoglycan biosynthesis pathway was positively correlated with the mean duration of grooming sessions in the Splash test, *r*_(14)_=0.657, *p*=.011, but negatively correlated with the number of grooming sessions in this test, *r*_(14)_=-0.804, *p*<.001. This pathway was also negatively correlated with the number of entries in the corners, *r*_(19)_=-0.682, *p*=.001, and the distance traveled, *r*_(19)_=-0.570, *p*=.011, in the open field. In males, only the peptidoglycan biosynthesis pathway was positively correlated with behavioral outputs, namely the mean duration of grooming sessions, *r*_(17)_=0.596, *p*=.012, in the Splash test.

**Figure 9:**
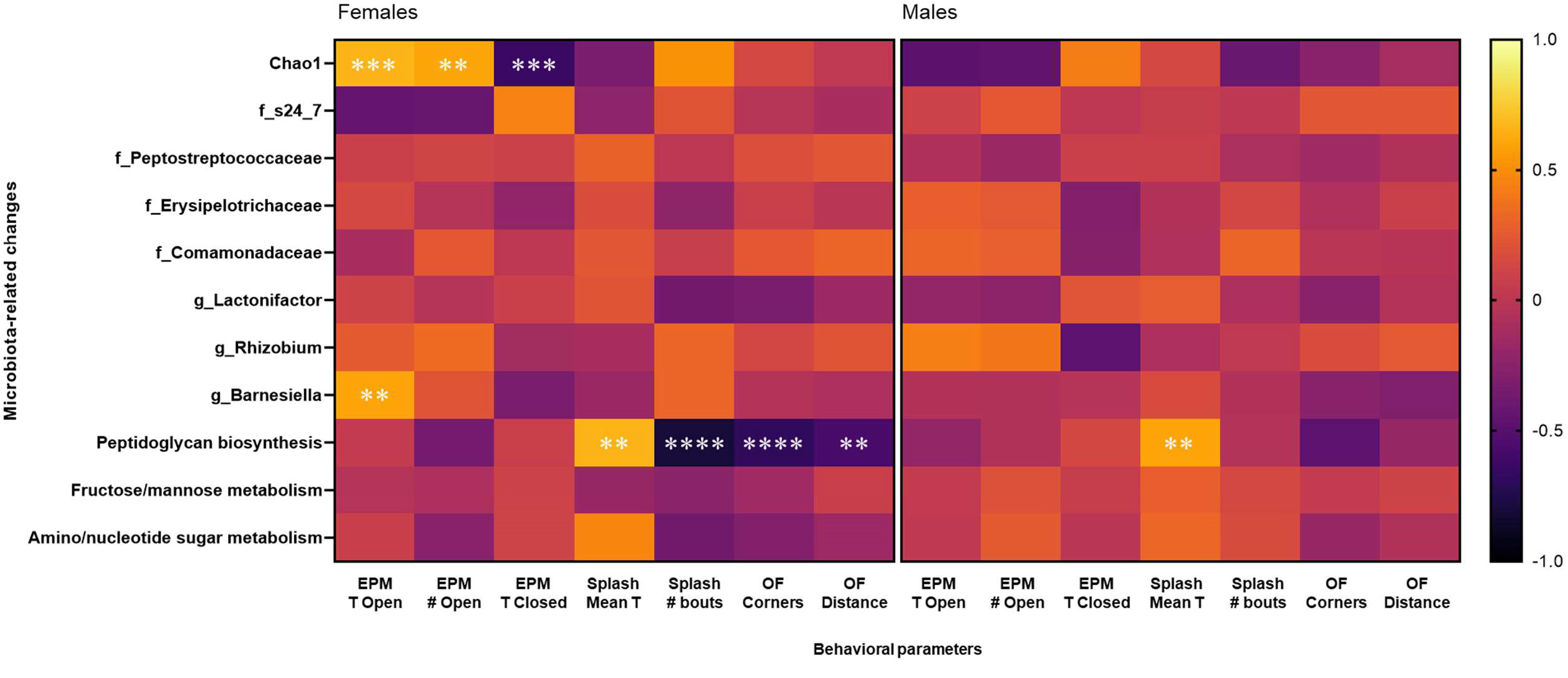
Relationships between microbiota-related outcomes and behavioral parameters affected by Exercise and/or Sex among female and male mice. In females (left panel), Chao1 positively correlated with the time spent (EPM T Open) and entries made into (EPM # Open) the open arms of the elevated plus maze but negatively correlated with the time spent in the closed arms (EPM T Closed) of the maze. The relative abundance of *Barnesiella* also positively correlated with the time spent in the open arms of the EPM. Enrichment of the peptidoglycan synthesis pathway positively correlated with the mean time spent in grooming bouts (Splash Mean T) in the Splash test but negatively correlated with the number of grooming bouts (Splash # bouts) in this test as well as with the number of entries in the corners (OF Corners) of and the distance traveled (OF Distance) in the open field. In males (right panel), the peptidoglycan synthesis pathway positively correlated with the mean time spent in grooming bouts in the Splash test. \*\**p*<.01, \*\*\**p*<.005, and \*\*\*\**p*<.001.

## 4. Discussion

Physical activity is being increasingly recognized as an effective approach to prevent and/or limit anxiety and depression, but the mechanisms underlying its mental health-improving effects are not yet entirely understood. As very few studies examining the psychological and physiological benefits of exercise in human populations have included sex as a biological variable, and as most studies conducted in rodent models have included only one sex, very little is known about the possibility that exercise could improve mental health differently among females and males. Here we show that short-term voluntary access to running wheels in adult C57BL/6N mice limited weight gains, attenuated behavioral parameters suggestive of anxiety, and changed grooming patterns exclusively in males. These effects appear independent of inflammatory changes in the intestinal and brain environments, at least in the regions examined in our study, as jejunal pro-inflammatory cytokine reductions in exercised mice were apparent only in females and hippocampal downregulation of the microglia-specific fractalkine receptor C3xcr1 occurred in both sexes. Interestingly, some of the behaviors assessed after the exercise regimen were linked to microbiota outcomes in a sex-dependent manner, suggesting that the behavioral status associated with anxiety- and depressive-like phenotypes in females versus males could be associated with the basal composition of their gut microbiota and/or with how gut microbial communities changed over the course of the exercise intervention.

Despite the total amounts of exercise being comparable among sexes, voluntary access to running wheels limited weight gains in males but not in females, suggesting that the metabolic response to exercise may have occurred differently in each sex. In line with this possibility, decreases in fat mass, improvements in mitochondrial function, and increases in adipokine secretion in white adipose tissue apparent in male mice with voluntary access to running wheels were absent in females (Nigro et al., 2021). We did not assess physiological metabolic parameters directly, but predictive microbiota functional outputs inferred from the PICRUSt2 analysis indicate that two pathways related to carbohydrate metabolism, the fructose and mannose metabolism and the amino sugar and nucleotide sugar metabolism, were enriched in males but not in females. Although further analyses will be needed to confirm which bacterial metabolites may be involved in this process, this raises the possibility that particular populations of bacteria could have contributed to body weight status during exercise in a sex-dependent way, through their actions on carbohydrate metabolism.

In addition to the male-specific effects of exercise on body weight, voluntary access to running wheels reduced fear in the anxiogenic parts of the elevated plus maze and changed grooming patterns in males only. The increased time spent in the open arms of the maze and the higher number of entries made into these arms could not be explained by an overall increase in activity in this test, as the distance traveled in the apparatus was comparable between groups. Reductions in anxiety-like behavior in the elevated plus maze have been reported in male mice subjected to different duration protocols of voluntary wheel running (Duman et al., 2008; Liśkiewicz et al., 2020; Santos-Soto et al., 2013). Similar effects have also been observed in females (Aujnarain et al., 2018), but in this particular study mice were housed in groups in an enriched environment that included a running wheel from weaning onward, which could have compounded the effects of structural and social enrichment with that of physical activity. In contrast, voluntary exercise did not have any fear-reducing effects in the open field but increased overall locomotor activity in both sexes. Opposing effects in these two tests after physical exercise have been reported by others (Binder et al., 2004), potentially owing to the residual effects of physical activity on locomotor activity in the open field, although the exact mechanisms responsible for this discrepancy remain to be determined.

The overall time mice spent grooming in the Splash test, which is the main indicator of motivation for self-care and suggestive of depressive-like behavior, was not changed in exercised mice of either sex. Yet, male mice with access to exercise initiated more but shorter grooming bouts. Similar to our findings, voluntary wheel running in adolescent male mice prevented decreases in the frequency of grooming elicited by repeated social defeat (Calpe-López et al., 2022), but to our knowledge no study has examined the effects of free running on grooming in otherwise basal conditions. We cannot exclude the possibility that the water mist administered to otherwise healthy mice, combined with their introduction to a novel environment, triggered low to moderate arousal. Arousing stimuli in rodents have been shown to elicit spontaneous grooming bouts that are more frequent but of shorter duration (Fernández-Teruel & Estanislau, 2016; Kalueff et al., 2016). In the absence of overall difference in the time invested in grooming, it is difficult to appreciate the effects of exercise on behaviors suggestive of depression, especially as parameters in the tail suspension test were unaffected by exercise, in line with previous reports (Calpe-López et al., 2022).

We next determined gene expression of pro-inflammatory factors in the hippocampus and in the jejunum. While Cx3cr1 expression in the hippocampus was downregulated by exercise in both sexes, the pronounced jejunal reductions of IL-6 and TNF-α expression after exercise were apparent in females only, indicating that inflammatory-related changes in the brain and intestinal environments after voluntary wheels running were not entirely coordinated. In the brain, Cx3cr1 is primarily expressed by microglia (Jung et al., 2000) and its exclusive ligand fractalkine (Cx3cl1) is predominantly secreted by neurons, thus allowing neurons to have a unique control over microglia activation and pro-inflammatory release (Hatori et al., 2002). The Cx3cr1 reductions in our exercised mice were not accompanied by changes in Iba1 or in pro-inflammatory cytokines, potentially owing to microglia being in a resting state, making them less responsive to the effects of exercise manipulations than microglia activated by aging or stress (He et al., 2017; Kohman et al., 2013; Xiao et al., 2021b). While this change in Cx3cr1 could be reflective of a disrupted neuron-microglia crosstalk, the functional consequences of this disruption are not entirely clear, especially as both protective and deleterious effects of Cx3cr1 deficiency have been reported (Pawelec et al., 2020; Sheridan & Murphy, 2013).

Voluntary wheel running has been shown to limit intestinal inflammatory activation in a mouse model of colitis using males (Cook et al., 2013) and to decrease TNF-α in lymphocytes from the small and large intestine in healthy female mice (Hoffman-Goetz et al., 2009). To our knowledge, we are the first to report sex-specific anti-inflammatory effects of exercise in the intestinal environment in the absence of an inflammatory challenge, although we cannot completely exclude the possibility that individual housing conditions promoted intestinal inflammatory activation, as previously reported in the bloodstream in response to individual housing (Du Preez et al., 2020; Tan et al., 2021). Why the cytokine-reducing effects of exercise in this part of the small intestine were specific to females is not entirely clear, but the finding suggests that free access to exercise modulates intestinal health in otherwise healthy individuals, at least in this sex, although further studies will be needed to determine whether this is the case in other parts of the digestive tract. In addition to the observation that exercise improves the quality of life in individuals with gastrointestinal inflammatory diseases (Klare et al., 2015), our findings support the view that short-term voluntary exercise may reduce the risk for these diseases, as previously suggested (Packer et al., 2010).

Unexpectedly, bacterial species richness was reduced in males with voluntary access to exercise, which contrasts with the accepted view that physical activity improves intestinal health through its effects on the diversification of gut bacterial communities (Clarke et al., 2014). Having a microbiota that was less rich was also associated with poorer outcomes in the elevated plus maze in females but, curiously, the apparent negative associations between microbiota richness and behavioral outcomes in males failed to reach significance. Although no overall effects of exercise or sex on microbiota composition (as determined by beta-diversity metrics) was observed, potentially due to the short-term duration of our exercise protocol (Allen et al., 2017), fluctuations at lower taxonomic ranks were detected. In accordance with previous reports (Evans et al., 2014), abundance of the butyrate-producing *S24-7* was increased in both female and male mice with access to running wheels, supporting a role for exercise in intestinal health improvements. Importantly, *Peptostreptococcaceae* and *Erysipelotrichceae* were changed by exercise in a sex-dependant manner, with *Peptostreptococcaceae* being reduced in males and *Erysipelotrichceae* being reduced in females. *Peptostreptococcaceae* has been reported to interact with host pathways involved in the regulation of intestinal homeostasis and inflammation in the context of intestinal inflammatory diseases (Priya et al., 2022). Similar to our findings, voluntary access to running wheels limited *Peptostreptococcaceae* elevations in male mice fed a diet rich in fat (Kang et al., 2014; Li et al., 2021) and diets enriched with phenolic compounds reduced basal *Peptostreptococcaceae* abundance in male mice, in combination with improvements of other markers of intestinal health (Monk et al., 2016). As for *Erysipelotrichaceae*, members from this family have been shown to have immunogenic properties (Palm et al., 2014), to correlate with markers of bacterial translocation and of systemic inflammation (Dinh et al., 2015), and to be elevated in obese individuals (Zhang et al., 2009). Notably, abundance of *Erysipelotrichaceae incertae sedis* was negatively correlated with depression severity in females (Chen et al., 2018). In our study, a decrease of *Erysipelotrichaceae* was apparent in females only, but reductions of species from this family have also been observed in male mice exposed to a longer duration regimen of free running (Choi et al., 2013). Why *Peptostreptococcaceae* and *Erysipelotrichaceae* in our exercised mice were reduced in a sex-dependent way has yet to be determined, but these changes support the view that exercise may contribute to intestinal health through its actions on bacterial populations that vary according to the sex of animals.

A very particular set of sex-specific associations between microbiota and behavioral outcomes were established. Although females had less *Barnesiella* than males, having more bacteria from this genus was associated with better outcomes in the elevated plus maze in this sex. *Barnesiella* have been found to protect the gastrointestinal tract from antibiotic-resistant strains (Ubeda et al., 2013) and to confer anti-inflammatory properties in the context of an inflammatory challenge (Weiss et al., 2014). The observation that higher *Barnesiella* abundance in females was associated with less behaviors suggestive of an anxiety-like phenotype may suggest that their limited presence in basal conditions, compared to males, could have interfered with the establishment of behavioral improvements throughout the exercise procedure. Similarly, the peptidoglycan biosynthesis pathway was unchanged in exercised males and impoverished in exercised females, but an enrichment in this pathway was associated with positive outcomes in the Splash test in both sexes as well as with reduced activity in the open field in females. Considering that microbiota-derived peptidoglycan can translocate to the bloodstream and reach the brain in mice under basal conditions, and that mice lacking the peptidoglycan-sensing protein Pglyrp2 had age- and sex-specific behavioral and locomotor impairments (Arentsen et al., 2017, 2018), our findings support a role for this bacterial motive in behaviors suggestive of mental health symptoms after exercise.

Females and males ran a comparable distance in their home-cage wheels, indicating that they performed a similar amount of exercise during the study. Apart from *Peptostreptococcaceae* and *Lactonifactor*, none of the outcomes determined in the current study were associated with the amounts of exercise, indicating that the body weight status and behavioral changes in males, the jejunal anti-inflammatory phenotype in females, and the remaining sex-specific microbiota fluctuations were independent of how much mice exercised throughout the duration of the experiment. In sum, our findings suggest a role for short-term voluntary exercise in the prevention of behaviors suggestive of anxiety and in intestinal health that is independent of exercise amounts, and that vary between females and males. Although intriguing, the observation that only males benefited from the behavioral effects of this exercise procedure suggests that other exercise regimen (e.g., longer duration or more structured) could be more suitable for females. It is also possible that behavioral tests different from the ones used in the current study could better capture mental health-related improvement related to voluntary exercise in females. On an exploratory note, we suggest that targeting *Barnesiella* and/or bacteria known to enrich the peptidoglycan biosynthesis pathway could represent a potential avenue in improving mental health symptoms, especially in females.

## Acknowledgments

This work was supported by the Natural Sciences and Engineering Research Council (NSERC #RGPIN-2016-06146 Discovery Grant to MCA). The NSERC Undergraduate Student Research Awards (to ZW), the Canadian Institutes for Health Research (Canada Graduate Scholarship – Master’s Program to NO and CN and Vanier PhD Canada Graduate Scholarship to JKS and MCU), the Ontario Graduate Scholarship Program (PhD Scholarship to AS), and Mitacs (Mitacs Research Training Award to ZW) also supported this research.

## Author Contributions

ZW and MCA designed the experiment. ZW, NO, BA, CN, and MCU conducted the experiment and performed the behavioral and molecular (RT-qPCR) analyses. ZW, NO, AS, and JKS conducted the microbiota sequencing analyses. JKS performed the bioinformatics analyses with the assistance of ZW. ZW and MCA interpreted the data and wrote the manuscript, which was edited by all authors.

---

**Supplementary Figure 1:**
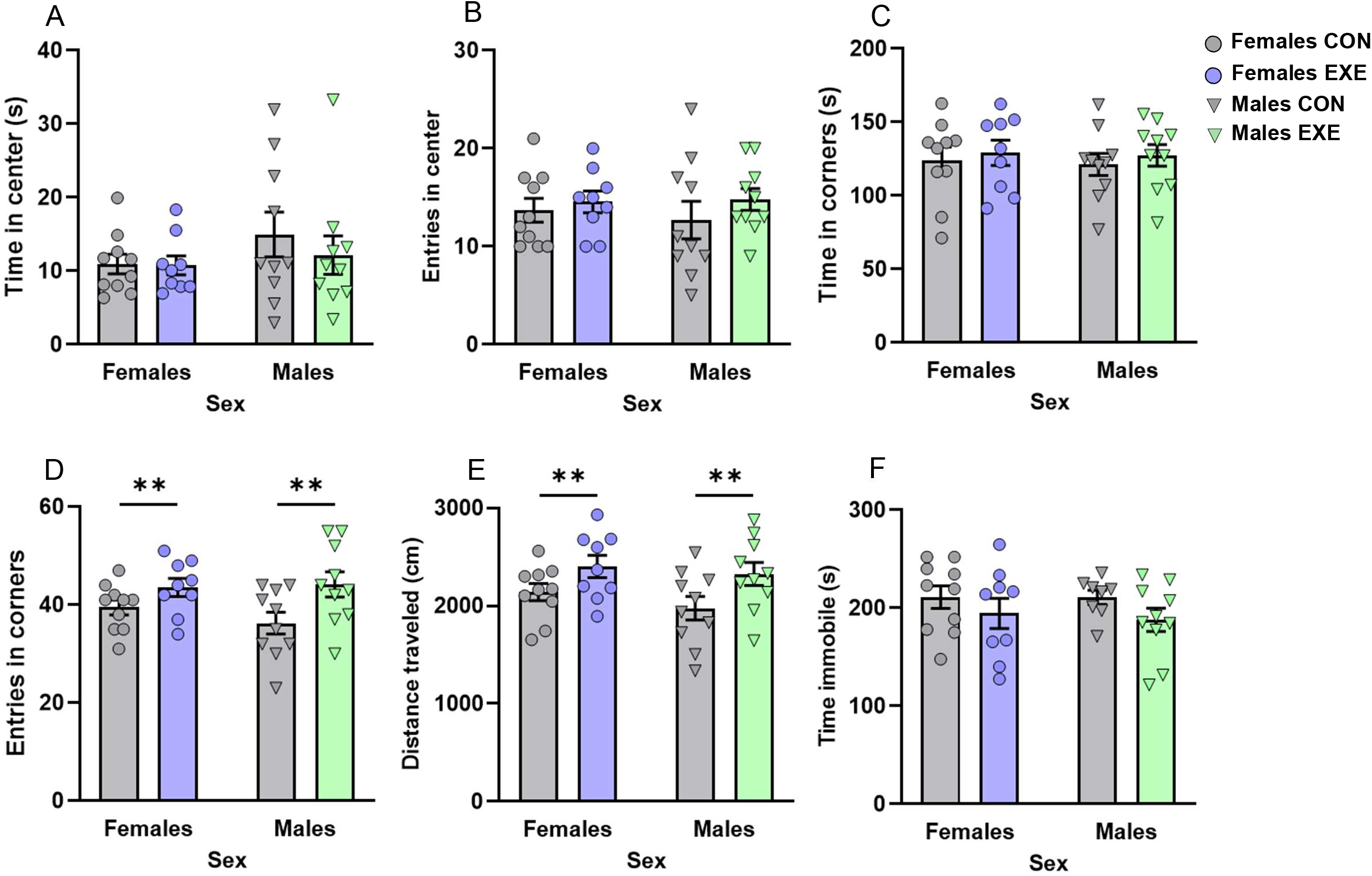
Behavioral parameters in the open field and tail suspension tests. (A) Time spent in the center (in seconds [s]), (B) number of entries into the center, and (C) time spent in the corners (in seconds [s]) of the open field are comparable among groups. Females and males with exercise (D) made more entries into the corners of the open field and (E) traveled more distance in the apparatus than their counterparts without exercise did. (F) Time spent immobile in the tail suspension test (in seconds [s]) was comparable among groups. Dots and triangles represent individual mice and bar plots and error bars represent group means ± S.E.M. Females without exercise (Females CON: *n*=10); females with exercise (Females EXE: *n*=9); males without exercise (Males CON: *n*=8-10); males with exercise (Males EXE: *n*=10). \*\**p*<.01 relative to sex-matched CON.

**Supplementary Figure 2:**
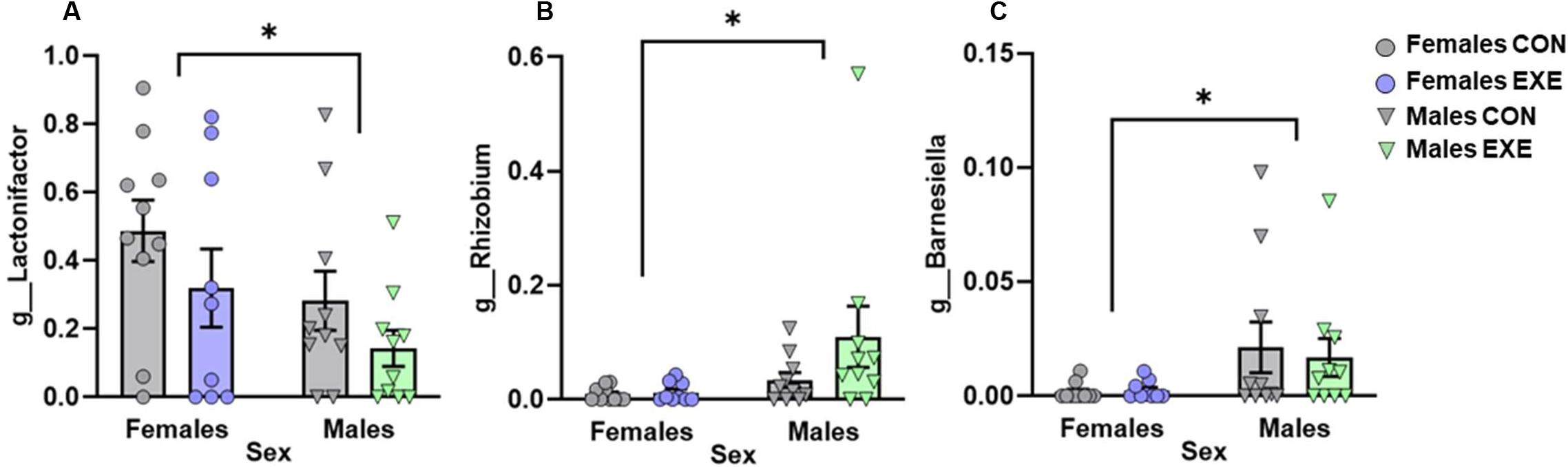
Changes in the relative abundance of genera based on sex only. (A) Relative abundance of the *Lactonifactor* genus was increased in females compared to males whereas that of the (B) *Rhizobium* and (C) *Barnesiella* genera was lower in females than in males. Dots and triangles represent individual mice and bar plots and error bars represent group means ± S.E.M. Females without exercise (Females CON: *n*=10); females with exercise (Females EXE: *n*=9); males without exercise (Males CON: *n*=10); males with exercise (Males EXE: *n*=10). \**p*<.05 relative to males.

**Supplementary Figure 3:**
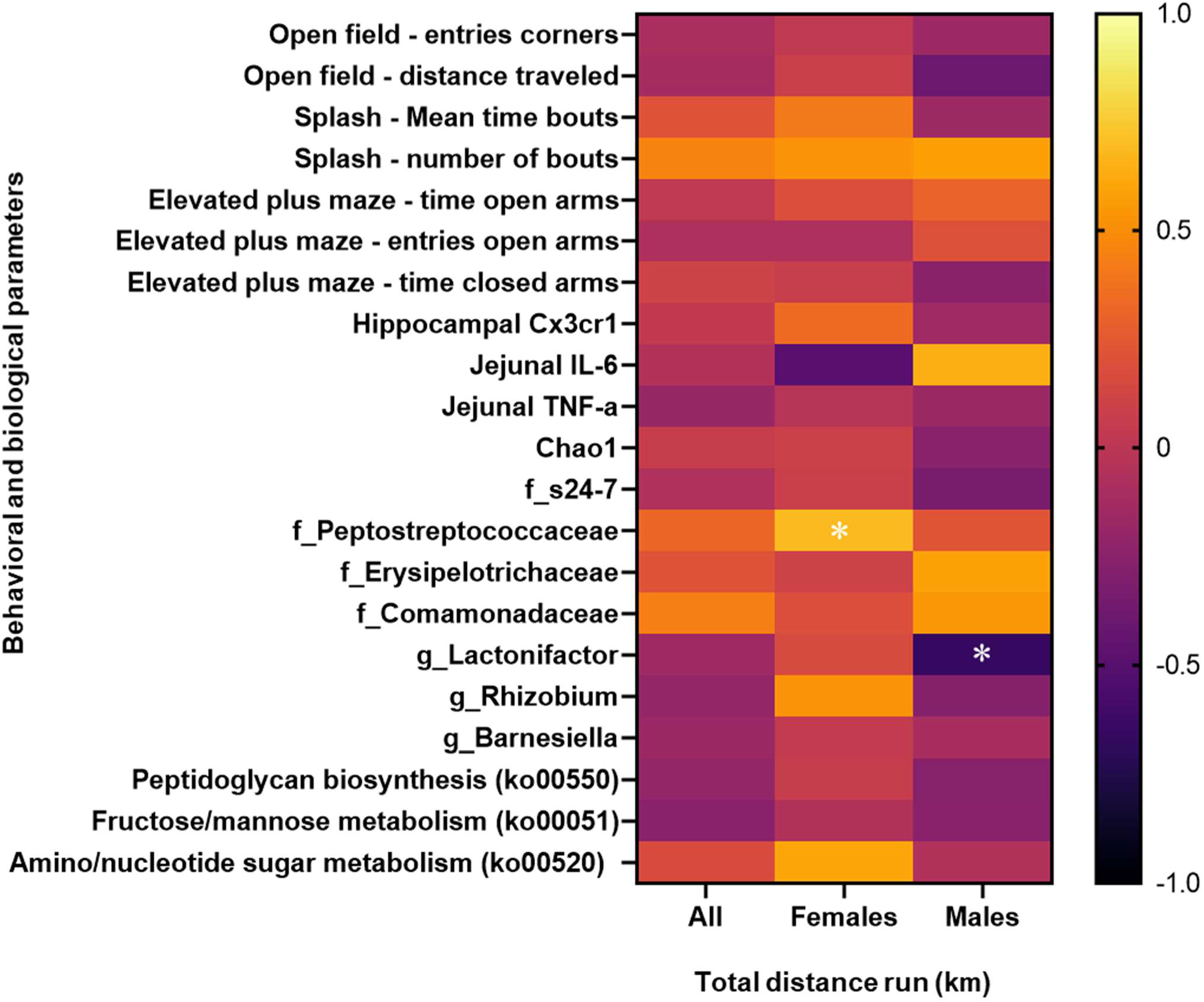
Relationships between behavioral and biological parameters affected by Exercise and/or Sex and the total distance run (in km) among female and male mice. Behaviors in the open field, Splash, and elevated plus maze tests as well as hippocampal and jejunal factors affected by either Sex and/or Exercise were not correlated with the total distance run (in km) in the wheels during the exercise regimen. The distance run in the wheels positively correlated with the relative abundance of the *Peptostreptococcaceae* family in females and negatively correlated with the relative abundance of the *Lactonifactor* genus in males. \**p*<.05.

**Supplementary Table 1.**
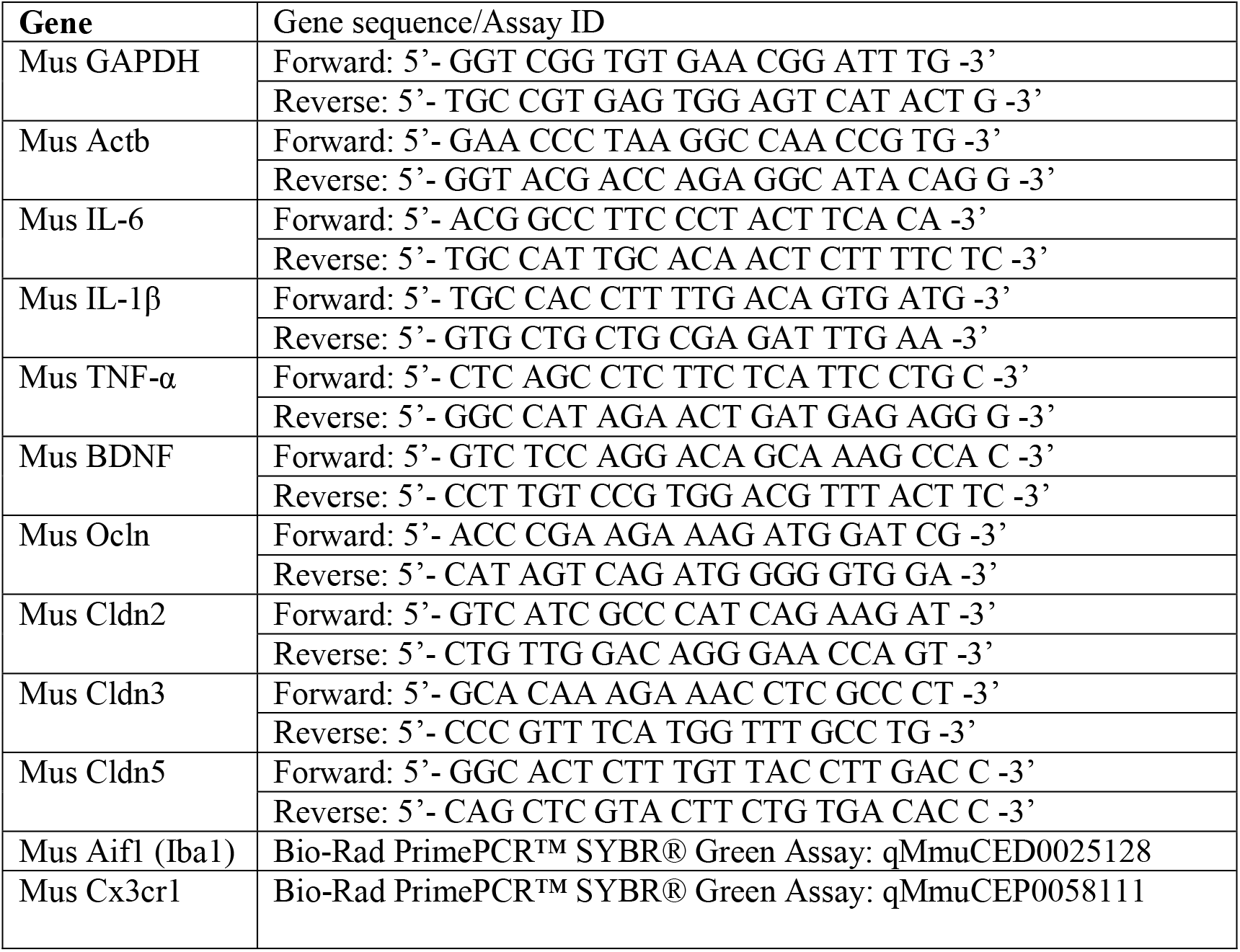
Primer sequences used in RT-qPCR analyses.

**Supplementary Table 2.** Gene expression of pro-inflammatory and tight junction markers in the hippocampus.

| Gene | Females |  | Males |  |
| --- | --- | --- | --- | --- |
|  | Controls | Exercise | Controls | Exercise |
| **IL-6** | 1.044 ± 0.192 | 1.045 ± 0.128 | 1.133 ± 0.174 | 1.049 ± 0.139 |
| **TNF-α** | 0.920 ± 0.116 | 1.028 ± 0.108 | 1.233 ± 0.128 | 0.979 ± 0.125 |
| **IL-1β** | 0.921 ± 0.182 | 1.228 ± 0.160 | 1.103 ± 0.207 | 0.877 ± 0.098 |
| **Ocln** | 0.886 ± 0.067 | 1.141 ± 0.119 | 1.033 ± 0.066 | 1.041 ± 0.112 |
| **Cldn5** | 1.139 ± 0.119 | 1.020 ± 0.112 | 0.912 ± 0.079 | 1.020 ± 0.136 |
| **Iba1** | 0.891 ± 0.090 | 1.181 ± 0.184 | 1.053 ± 0.085 | 1.001 ± 0.081 |

